# Triggering receptor expressed on myeloid cells 2 (TREM2) regulates phagocytosis in glioblastoma

**DOI:** 10.1101/2023.04.05.535792

**Authors:** Mekenzie M. Peshoff, Pravesh Gupta, Rakesh Trivedi, Shivangi Oberai, Prashanth Chakrapani, Minghao Dang, Nancy Milam, Mark E. Maynard, Brian D. Vaillant, Jason T. Huse, Linghua Wang, Karen Clise-Dwyer, Krishna P. Bhat

## Abstract

Glioblastomas (GBMs) are tumors of the central nervous system that remain recalcitrant to both standard of care chemo-radiation and immunotherapies. Emerging approaches to treat GBMs include depletion or re-education of innate immune cells including microglia (MG) and macrophages (MACs). Here we show myeloid cell restricted expression of triggering receptor expressed on myeloid cells 2 (TREM2) across low- and high-grade human gliomas. TREM2 expression did not correlate with immunosuppressive pathways, but rather showed strong positive association with phagocytosis markers such as lysozyme (LYZ) and CD163 in gliomas. In line with these observations in patient tumors, *Trem2^-/-^* mice did not exhibit improved survival compared to wildtype (WT) mice when implanted with mouse glioma cell lines, unlike observations previously seen in peripheral tumor models. Gene expression profiling revealed pathways related to inflammation, adaptive immunity, and autophagy that were significantly downregulated in tumors from *Trem2^-/-^* mice compared to WT tumors. Using ZsGreen-expressing CT-2A orthotopic implants, we found higher tumor antigen engulfment in Trem2^+^ MACs, MG, and dendritic cells. Our data uncover TREM2 as an important immunomodulator in gliomas and inducing TREM2 mediated phagocytosis can be a potential immunotherapeutic strategy for brain tumors.

**Key points:**

- TREM2 is not associated with immunosuppressive molecules in GBM
- TREM2 is associated with phagocytosis in both human and mouse gliomas
- Deletion of Trem2 in mice does not improve survival in glioma models

**Importance of the study:** Triggering receptor expressed on myeloid cells 2 (TREM2) has been implicated as a major immunoregulator in both neurodegenerative diseases and systemic cancers, yet its functional role in gliomas remains unclear. This study reveals that unlike in other cancers, TREM2 is not associated with immunosuppression in the glioma microenvironment. In fact, TREM2 expression is associated with phagocytosis in both human and mouse gliomas, similar to its role in Alzheimer’s disease. These findings indicate that TREM2 blockade will not be a viable treatment strategy for gliomas. Instead, TREM2 induction may boost the potential of myeloid cells in the tumor microenvironment to engulf cancer cells.

## Introduction

Gliomas are the most common primary adult brain tumors and comprise over 80% of central nervous system (CNS) malignancies (Ostrom, et al., 2015). Of these, glioblastomas (GBMs) are high grade gliomas that exhibit the poorest clinical outcome. Despite standard of care treatments including surgical resection and concurrent chemoradiation, these patients have a median survival of only 15-18 months (Stupp et al., 2005) (Hegi et al., 2005). Although lymphocyte-based immune checkpoint therapies have significantly improved progression-free survival for many other solid tumors, these modalities have yielded disappointing results in clinical trials for GBM patients (Reardon et al., 2020) (Sade-Feldman et al., 2018). In fact, glioma patients are lymphopenic (Kim et al., 2019) and inherently show T cell scarcity in the tumor immune microenvironment (TIME, Klemm et al., 2020) (Friebel et al., 2020), yet currently approved immunotherapies target T lymphocytes for facilitating antitumor immunity.

Recent immunophenotyping studies have revealed that the glioma TIME is characterized by a heterogeneous, predominantly myeloid population (Chen et al., 2017) (Larkin et al., 2022) (Gupta et al., 2022 *Preprint*). These myeloid cells include yolk sac-derived brain resident microglia (MG), bone marrow-derived macrophages (MACs), and monocyte-derived macrophages (MDMs) (Morantz et al., 1979) (Gupta et al., 2022 *Preprint)*. Because of their relative abundance, selective targeting of myeloid cells may be a more viable alternative to current T cell-based investigations (Quail and Joyce, 2017). Myeloid-targeted clinical trials in various cancers are emerging (Cassetta and Pollard, 2018), but a deeper functional understanding of anti-tumor properties of myeloid cells in GBM is needed. Although MG and MACs perform a variety of protective immunoregulatory functions including phagocytosis of pathogens and cellular debris as well as antigen presentation to T cells (Barker et al., 2002) (Schetters et al., 2017) (Brown and Neher, 2014), how these pleiotropic functions affect glioma progression remains unclear.

Triggering receptor expressed on myeloid cells 2 (TREM2) is a type I transmembrane receptor in the immunoglobulin superfamily that has been implicated as a major regulator of the myeloid cell immune response (Bouchon et al., 2001) (Deczkowska et al., 2020). It is expressed on bone marrow-derived MACs, MG, and is implicated in dendritic cell (DC) maturation (Bouchon et al., 2001). The binding of its various ligands and association with DNAX activator proteins 10 and 12 (DAP10, DAP12) result in activation of the PI3K-Akt pathway or SYK kinase, resulting in downstream promotion of cell survival pathways or phagocytosis and cytokine production, respectively (Peng et al., 2010). TREM2 is involved in engulfment, a crucial step for the initiation of phagocytosis (Takahashi et al., 2005), and TREM2 deficient or defective MG display impaired phagocytosis *in vitro* and *in vivo* (Kleinberger et al., 2017) (N’Diaye et al., 2009) (Ulrich et al., 2014). TREM2 is well characterized in Alzheimer’s disease (AD), where the R47H mutation is a variant associated with an increased risk of typical early-onset AD (Jonsson et al., 2013) (Guerreiro et al., 2013) and TREM2 promotes microglial survival and phagocytosis of amyloid plaques in early-stage AD mouse models (Wang et al., 2015) (Ulland et al., 2017) (Wang et al., 2016). Recently, TREM2 has garnered attention for its role in cancer due to its expression on tumor associated MACs and an increasing interest in the dual role of myeloid cells in the inflammatory response against tumors as well as pro-tumoral immunosuppression (Molgora et al., 2020) (Katzenelenbogen et al., 2020). Despite its critical role in other cancers and neurological disorders, the function of TREM2 in gliomas has not been well-characterized, and previous studies lack *in vivo* relevance and provide only correlative associations (Yu et al., 2022).

In this study, we characterize the role of TREM2^+^ myeloid cells using both human patient-derived glioma tissues as well as orthotopic mouse models. Through a comprehensive analysis of a large cohort of human gliomas, we identified a population of glioma-associated TREM2^+^ myeloid cell populations that are associated with phagocytosis. We also find that Trem2^+^ myeloid cells engulf tumor antigens, and survival benefits of Trem2 deletion as reported in other cancers (Molgora et al., 2020) are not observed in mouse models of glioma. In summary, our pre-clinical and translational studies define TREM2 as a key phagocytic immunomodulator in gliomas.

## Methods

### Human brain tissue collection

Tissue samples were collected from fifty-six glioma patients and five epilepsy patients. Informed consent and detailed information including age, sex, glioma type, and site of tumor extraction were obtained prior to neurosurgery. The brain tumor samples were collected according to MD Anderson internal review board (IRB)-approved protocols LAB03-0687, LAB04-0001, and 2012-0441, and epileptic brain tissue was collected at Baylor College of Medicine under IRB-approved protocol H-13798. All experiments performed were in compliance with the IRB of MD Anderson Cancer Center.

### Preparation of single-cell leukocyte suspensions

Resected brain tumors or tissue from humans and mice were freshly processed or stored overnight in MACS tissue storage solution (Miltenyi Biotech #130-100-008). Human tissue and mouse lymph nodes and spleen were finely minced, and mouse brains were homogenized using a Dounce homogenizer. Tissues were enzymatically dissociated in warm digestion medium containing 100 µg/mL collagenase D (Sigma-Aldrich #11088866001) and 2 µg/mL DNase for 30 minutes at 37° C. The reaction was neutralized with 2% fetal bovine serum (FBS) (Gibco #16140-071) in IMDM. Tissues were passed through a 100 µm cell strainer (Corning #352360) and brain samples were subjected to 33-40% Percoll gradient (Sigma-Aldrich #17-0891-01) to remove myelin then resuspended in red blood cell (RBC) lysis buffer (Sigma-Aldrich #R7757-100ML) for 10 minutes at room temperature. The RBC lysis reaction was neutralized with an equal volume of 1X PBS. Cell pellets were obtained via centrifugation at 500g for 5 minutes at 4° C, and human samples were cryopreserved in 10% DMSO (Sigma-Aldrich #F4135) in FBS in liquid nitrogen at -196°C until use, whereas mouse tissues were collected and processed freshly ahead of experimentation. A detailed description of methods for human leukocyte preparation is provided in Gupta, et al., 2022.

### IHC and immunofluorescence

Formalin fixed paraffin embedded (FFPE) tissues from human patients and mice were serially sectioned on a microtome at 5 µM. Slides were baked for 30 minutes at 60° C then deparaffinized by subsequent washes in xylene, 1:1 xylene and 100% ethanol, 100% ethanol, 95% ethanol, 70% ethanol, 50% ethanol, and water. Antigen retrieval was performed using a pressure cooker at 100°C for 10 minutes in sodium citrate pH=6 buffer. Slides were washed 3 times in 1X PBS and endogenous peroxidases were blocked with 3% hydrogen peroxide for 30 minutes. After 3 washes in PBS, slides were incubated for an hour at room temperature with 10% bovine serum albumin (Millipore Sigma #A7030) to block nonspecific binding. Slides were incubated overnight at 4°C with primary antibody (1:100 TREM2 (D8I4C) rabbit mAb, Cell Signaling Technology #91086). The following day, after 3 washes in PBS, the slides were incubated with the corresponding HRPconjugated secondary antibody for one hour at room temperature (goat anti-rabbit IgG H&L HRP, Abcam #ab6721). The slides were then washed in PBS 3 times then developed using the DAB substrate kit (Abcam #ab64238). The slides were covered in hematoxylin for 3 minutes, then counterstained in PBS for 5-10 minutes. The slides were dehydrated and cover slipped using aqueous mounting medium. Quantitative analysis was performed blinded using ImageJ IHC toolkit by multiple authors.

### Gene module correlation analyses

For the correlation of phagocytic gene modules to the TREM2 pathway, we included ‘*TREM2’, ‘HCST’, ‘TYROBP’, ‘APOE’, ‘CLU’, ‘CD33’, ‘LGALS1’, ‘LGALS3’, ‘GRN’, ‘NFATC1’, ‘MS4A4A’, ‘MS4A6A’*. Phagocytic genes were curated from related gene ontology terms containing ‘Phagocytosis’ and the expression values are enlisted in Supplementary Table S1. TREM2 pathway score and phagocytic scores were calculated for each myeloid cell using Seurat function AddModuleScore (Butler et al., 2018).

### Animals

WT C57BL/6J (Jackson Labs #000664) and C57BL/6J-Trem2^em2Adiuj^/J (Jackson Labs #027197) mice were procured from Jackson Laboratory and housed at MD Anderson Department of Veterinary Medicine and Surgery under IACUC-approved protocol #00001504-RN02. Mice used for experiments were sex and age matched.

### Cell lines and *in vivo* glioma models

6-month-old mice were intracranially bolted at 2.5 mm lateral and 1 mm anterior to the bregma according to guide-screw procedure (Lal et al., 2000) One week following bolting, mice were implanted via stereotaxic injection with 20,000 CT-2A-luc, 20,000 CT-2A, or 100,000 GL261-luc cells resuspended in serum-free DMEM. For flow cytometry analysis, mice were implanted with 700,000 CT-2A-ZsGreen cells and sacrificed 12 days postinjection. Mice in all survival studies were monitored daily for signs of distress and were sacrificed when moribund, as defined by hunched posture, ataxia, and neurological sequelae including seizures. To monitor tumor growth via bioluminescence, mice were injected subcutaneously with 20 µL/g 15 mg/mL sterile D-luciferin and were imaged using the Perkin-Elmer in vivo imaging system (IVIS 200) 15 minutes post-injection. Data were analyzed using Aura (Spectral Instruments Imaging) software. CT-2A cells were a gift from Dr. Sourav Ghosh (Yale University) and fingerprinted prior to use. CT-2A-luc cells were purchased from Sigma-Aldrich (#SCC195) and expanded in-house. CT-2AZsGreen cells were generated using lentiviral particles derived from a pLVX-ZsGreen1C1 vector (Takara Bio #632566). All cell lines were grown in DMEM high glucose with 10% FBS (Gibco #16140-071) and CT-2A-ZsGreen cells were grown on a puromycin (Thermo Fisher #A1113803) selection cassette at 1 µg/mL. Cells were passaged twice a week or when 70-80% confluent. Culture medium was aspirated and flasks were washed with sterile 1X PBS, then cells were dissociated through trypsinization (Gibco #T3924100) for 3 minutes at 37°C followed by neutralization with an equal volume of culture medium. Cells were pelleted via centrifugation for 3 minutes at 400g, then resuspended in culture medium and split 2:1. All cells for experiments were used within the first 5 passages after thawing.

### RNA isolation and NanoString analysis

1.5 mm punch biopsy needles were used to extract tissue from the tumor core of FFPE mouse brains (Integra #33-31A). The QIAGEN RNeasy FFPE kit (QIAGEN #73504) was used to isolate RNA following deparaffinization. The RNA was sent for NanoString nCounter analysis using the mouse neuroinflammation panel. Statistical analysis of gene expression was performed according to NanoString protocol in the nSolver advanced analysis system and in R. Supplementary Table S2 shows the list of genes and their expression values in WT and Trem2^-/-^ murine tumors.

### Spectral flow cytometry and staining

Thawed cryopreserved human specimens were washed in 10% FBS in IMDM, pelleted at 500g for 5 minutes, and resuspended in 10% FBS in IMDM and incubated for 30 minutes at 37°C. Mouse cells were suspended in 10% FBS in IMDM upon dissociation. Cells were washed twice with 1X PBS then stained with Live Dead Blue Dead dye (Invitrogen #L34962) for 15 minutes at 4°C. Cells were washed three times in 1x PBS then stained for 15 minutes at 4°C in Fc block cocktail (1:20 Nova block, Phitonex, Mouse TruStain FcX, Biolegend #101320, CellBlox monocyte/MAC blocking buffer, Invitrogen #B001T07F01). Cells were stained in an antibody cocktail containing the appropriately titrated antibodies described in Supplementary Table S3 for 20 minutes at 4°C. Cells were washed in FACS buffer three times and fixed overnight in 200 µL True Nuclear fixation buffer (Biolegend #73158, #71360) at 4°C. Data were acquired using the Cytek Aurora 5 spectral flow cytometer. Data were analyzed using Cytek SpectroFlo and Becton Dickinson FlowJo 10.8.1.

### TCGA and GLASS cohorts

528 GBMs and 10 nontumor brains from the TCGA dataset analyzed on HG-U133A were used for all GBM correlation analyses (Cancer Genome Atlas Research, 2008). For LGG (grade I and grade II gliomas), the TCGA dataset containing 513 LGGs was analyzed (Cancer Genome Atlas Research et al., 2015), and pooled glioma analysis was performed using the TCGA GBMLGG combined set (Ceccarelli et al., 2016). Data were visualized and all analyses were computed using the GlioVis platform (gliovis.bioinfo.cnio.es) (Bowman et al., 2017). Data were generated by the TCGA Research Network (https://www.cancer.gov/tcga). Immunohistochemistry was performed on gliomas from the GLASS (Glioma Longitudinal AnalySiS) consortium (Consortium, 2018).

## Results and discussion

### Characterization of TREM2 expression in glioma-associated myeloid cells

Although TREM2 plays a central immunomodulatory role in neurodegenerative diseases, its role in gliomas has not been elucidated. Therefore, we evaluated *TREM2* mRNA expression across glioma associated myeloid and lymphoid cells. We used our single cell RNA sequencing human glioma dataset that provides the most advanced picture of immune contexture of isocitrate dehydrogenase (IDH) classified gliomas (Gupta et al., 2022 *Preprint)*. Myeloid cells showed significant enrichment of *TREM2* when compared to lymphoid cells across gliomas classified by IDH mutation status (Fig. 1A). These observations are consistent with previous studies demonstrating TREM2 restriction to myeloid lineage cells (Bouchon et al., 2001) (Qiu et al., 2021) and a recent study showing TREM2 expression in glioma-associated MG and MACs (Yu et al., 2022). We further interrogated *TREM2* expression in our transcriptionally defined myeloid cell types, which revealed its expression in MG, MG-like, MAC, Myeloid-IFN and cDC2 subsets compared to monocytes, neutrophils and other DCs (Fig. 1B). Immunofluorescence co-staining with MG/MAC marker Iba1 (Sasaki et al., 2001) further validated our genomic observations that TREM2 is tightly correlated with myeloid cells (Fig S1A and S1B). Because MG and MACs can be altered by treatment paradigms (Wei et al., 2020) (Gabrusiewicz et al., 2016), we evaluated TREM2 in matched treatment naïve versus recurrent gliomas from the GLASS (Glioma Longitudinal AnalySiS) consortium cohort (Consortium, 2018) using immunohistochemistry (IHC). When compared to IDH mutant (IDH-mut) gliomas, which displayed ramified morphology indicative of sensing (Nimmerjahn et al., 2005), TREM2^+^ cells showed predominantly amoeboid or activated morphology in IDH-wildtype (IDH-wt) gliomas (Fig. 1C). Furthermore, the proportion of TREM2^+^ cells in the tumor core were significantly higher in IDH-wt than IDH-mut gliomas (Fig. 1D), however no differences between primary and recurrent tumors were observed. These results suggest that differential infiltration of TREM2^+^ cells correlates with IDH status but is not necessarily altered by chemo-radiation.

**Figure 1.**
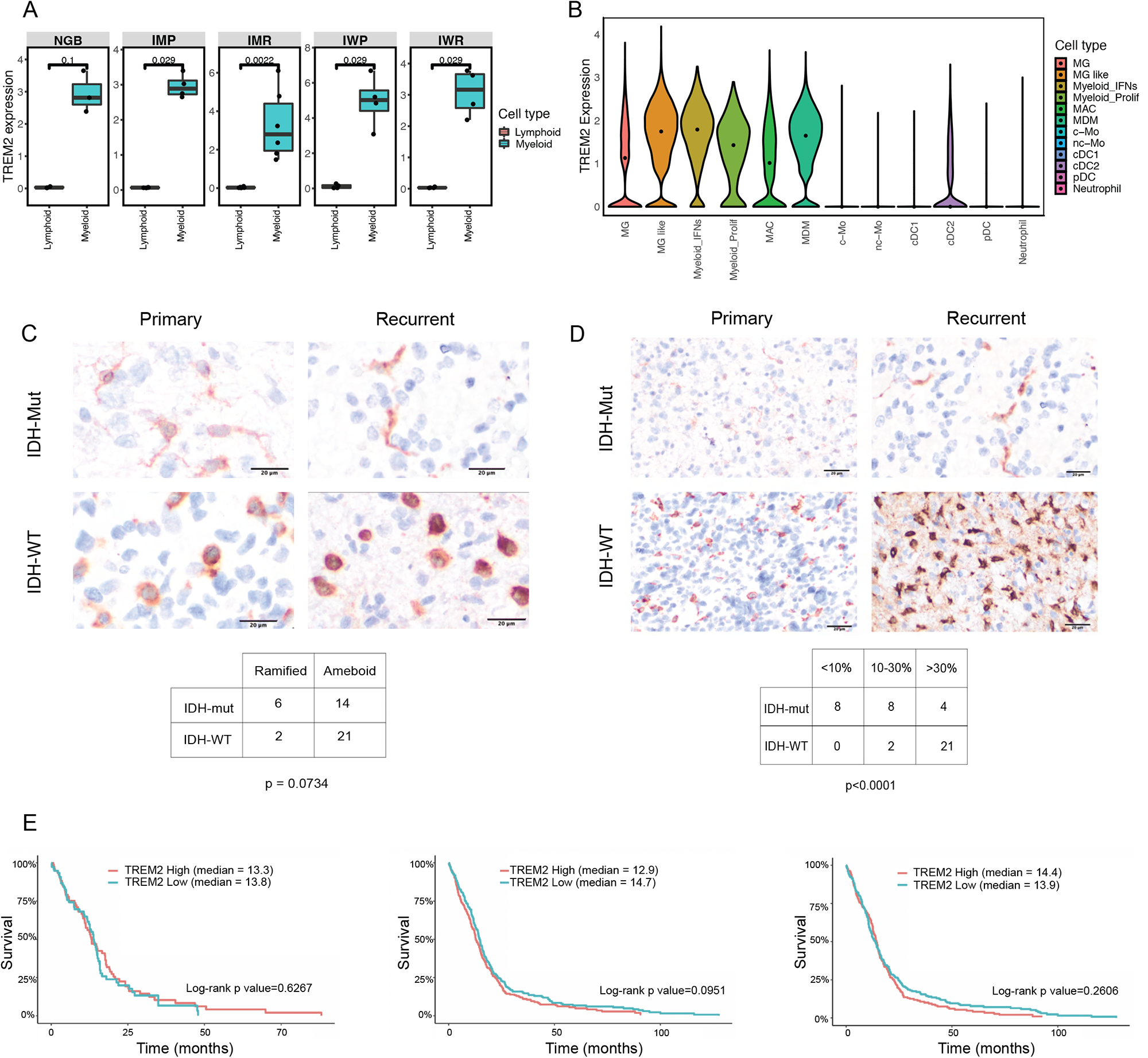
TREM2 is associated with myeloid cells and highly expressed in IDH-wt GBMs. **(A)** Bar graphs showing *TREM2* mRNA expression levels on lymphoid vs. myeloid cells across glioma types. NGB=non glioma brain, IMP=IDH-mut primary, IMR=IDH-mut recurrent, IWP=IDH-wt primary, IWR=IDH-wt recurrent. Wilcoxon signed-rank test was used to derive p values. **(B)** Violin plot of *TREM2* expression in myeloid cell subsets as defined by Gupta, et al. **(C)** Representative staining of TREM2 IHC in primary and recurrent IDH-wt and IDH-mut gliomas. Scale bar: 20 µm. Table shows TREM2^+^ cells were classified as ramified or ameboid based on extent of branching. p value was calculated with Chi-square goodness of fit. **(D)** Representative IHC images of TREM2 in primary and recurrent IDH-wt and IDH-mut tumors. Scale bar: 20 µm. The contingency table shows the number of gliomas with <10%, 10-30%, or >30% TREM2^+^ cells in the tumor core. p value was calculated using Chi-square goodness of fit. **(E)** Kaplan-Meier curves from TCGA GBM datasets (analyzed via RNA-Seq, HG-U133A, and Agilent-4502A, respectively) showing no significant survival differences in TREM2 high vs. TREM2 low GBMs. For RNA-Seq, log-rank p value=0.6267 and Wilcoxon p value=0.83553. For HG-U133A, log-rank p value=0.0951 and Wilcoxon p value=0.1032. For Agilent-4502A, log-rank p value=0.2606 and Wilcoxon p value=0.8442.

We extended these observations in a larger cohort of TCGA datasets and found that *TREM2* and its signaling partner *TYROBP* are associated with higher grade, which are predominantly IDH-wt tumors (Fig. S1C, S1D) and *TREM2* expression correlated with poor survival in low grade gliomas (LGG, Fig. S1E). However, *TREM2*-high versus -low GBMs displayed no significant survival differences (Fig. 1E). Based on the unique morphologies displayed by TREM2^+^ myeloid cells in our IHC cohort, these findings suggest that the role of TREM2 in the glioma microenvironment may be dependent on tumor grade and IDH status. Taken together, our genomic and proteomic analyses validated by larger datasets indicate that TREM2 is highly expressed in myeloid cell lineages in IDH-wt high grade gliomas.

### TREM2 correlates with microglial phagocytosis pathway molecules

Recent studies have shown that *TREM2* correlates with expression of immunosuppressive markers including *ARG1, MARCO, MRC1,* and *ITGA4* in peripheral malignancies such as sarcomas, breast, and colorectal cancers (Katzenelenbogen et al., 2020; Khantakova et al., 2022; Molgora et al., 2020). TREM2 blockade or deficiency in mice causes remodeling of the TIME that is associated with decreased tumor growth, alteration of MACs to express immunostimulatory molecules and increased infiltration of T lymphocytes and NK cells (Molgora et al., 2020). Our analyses of TCGA datasets showed that although TREM2 correlated with canonical pathway genes such as *APOE, TYROBP,* and *CD33* (Deczkowska et al., 2020), no correlation between *TREM2* and immunosuppressive genes were observed in either LGG or GBMs (Fig. 2A, S2A). TREM2 is known to be a phagocytic mediator, positively regulating the microglial ability to recognize and engulf pathogens through activation of its downstream pathways (Krasemann et al., 2017) (Yeh et al., 2016). We therefore examined if expression of TREM2 correlates with phagocytosis gene modules. We used a condensed metagene score derived from canonical genes participating in phagocytic functions including genes associated with phagosomes, vesicle mediated transport, and lysosomes (see methods and Supplementary Table S1). This analysis revealed that TREM2 expression significantly showed positive correlations with the phagocytic score primarily in MG compared to MAC and MDMs (Fig. 2B). We next examined TREM2 expression across different myeloid cell subsets by spectral cytometry using lysozyme (LYZ), a marker for active phagocytosis (Venezie et al., 1995) and the MAC scavenger receptor CD163, which plays a role in rendering MACs phagocytosis competent (Schulz et al., 2019) (Colton and Wilcock, 2010). TREM2 was confirmed to be expressed by both MG and non-MG myeloid cell populations in gliomas (Fig. 2C). We used the median fluorescence intensity (MFI) of LYZ and CD163 to define their distribution in TREM2 stratified myeloid subsets from primary and recurrent IDH-mut and IDH-wt tumors. Across all glioma subtypes, TREM2 was associated with higher expression of LYZ and CD163 in both microglial and MAC cell populations (Fig. 2D). In the TCGA GBM transcriptomic datasets, we found similar positive correlations between *TREM2* and *CD163*, *LYZ* and *CD74*, a gene encoding class II major histocompatibility complex (Basha et al., 2012) (Fig. 2E). These data demonstrate that TREM2 mRNA and protein correlate with phagocytosis markers in human gliomas.

**Figure 2.**
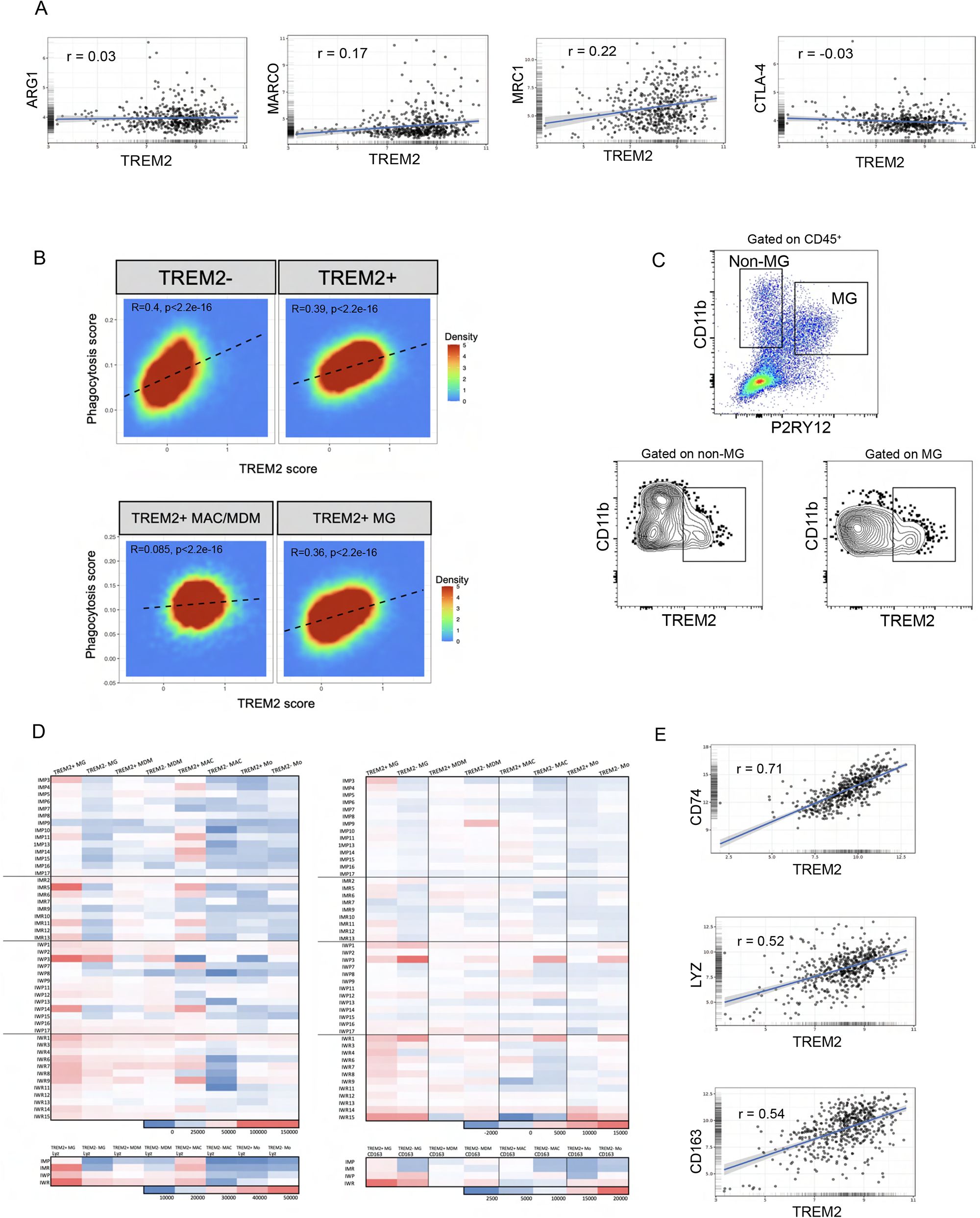
TREM2 is associated with phagocytosis in gliomas. **(A)** Linear regression plots showing lack of correlation between *TREM2* and immunosuppressive markers from TCGA GBM HG-U133A. r value was computed using Pearson’s product moment correlation. For all plots, p<0.001. **(B)** 2D density plot showing distribution of TREM2-and TREM2^+^ MG, MDMs, and MACs based on their phagocytosis and TREM2 pathway score. **(C)** Gating strategy showing TREM2 expression on both MG and non-MG myeloid subsets. MG are defined as CD11b^+^ P2RY12^+^, and non-MG myeloid cells are CD11b^+^ P2RY12-. In both MG and non-MG subsets, TREM2^-^ and TREM2^+^ populations are observed. **(D)** Heat map of median fluorescence intensity (MFI) values of LYZ and CD163 in TREM2^+^ and TREM2^-^ sorted myeloid populations across gliomas. Each row in the heatmap is data from a single patient. **(E)** Linear regression plots showing correlation of TREM2 expression with CD74, LYZ, and CD163 in TCGA GBM dataset HG-U133A. The r value was computed using Pearson’s product moment correlation. For all correlations, p<0.001.

### TREM2 expression on myeloid cells is dispensable for glioma growth

The role of MG and MACs in GBM progression is still evolving given their heterogeneity and lack of appropriate mouse models to completely demarcate these cell types. Nevertheless, previous studies have shown functional distinction of MG and MACs in gliomas. For example, MG regulate phagocytosis even in the absence of MACs (Hutter et al., 2019) whereas depletion of MACs restrains tumor progression (Chen et al., 2017; Pyonteck et al., 2013). Because TREM2 can be expressed by both myeloid populations, we sought to examine the impact of *Trem2* deletion on the growth of brain tumors. We initially used GL261, a widely used murine glioma line, transduced with luciferase (GL261-luc) and performed intracranial orthotopic implants in C57BL/6 mice. When compared to wild-type (WT) mice, *Trem2^-/-^* mice showed no significant differences in survival rates, which is in contrast to previous studies (Molgora et al., 2020) (Timperi et al., 2022) where *Trem2* deletion caused reduced tumor progression in sarcoma and triple-negative breast cancer xenografts (Fig. S2A). Gross histological examination of the tumors did not reveal differences in grade or morphology (not shown). Although the survival differences did not reach statistical significance, the kinetics of the tumor growth showed significant differences between WT and *Trem2*^-/-^ mice (Fig. S2B). Previous studies have shown that murine glioma models show differences in immune response that can be further altered by luciferase transduction (Podetz-Pedersen et al., 2014; Sanchez et al., 2020). To address this, we used syngeneic mouse glioma cell lines CT-2A and CT-2A-luc to assess the impact of *Trem2* depletion. Consistent with the trend observed with GL261, the *Trem2^-/-^* mice showed no significant differences in survival compared to WT mice (Fig. 3A, and 3B). Similarly, the kinetics of the tumor growth were significantly higher in *Trem2^-/-^* mice (Fig. 3C and 3D). These data imply that TREM2 is dispensable for tumor growth and in fact may play a tumor suppressive role in gliomas compared to systemic cancers. Further studies remain to be seen if the differences are due to differential roles and dynamics of Trem2^+^ MG versus peripheral MACs using cell type specific *Trem2* genetic ablation.

**Figure 3.**
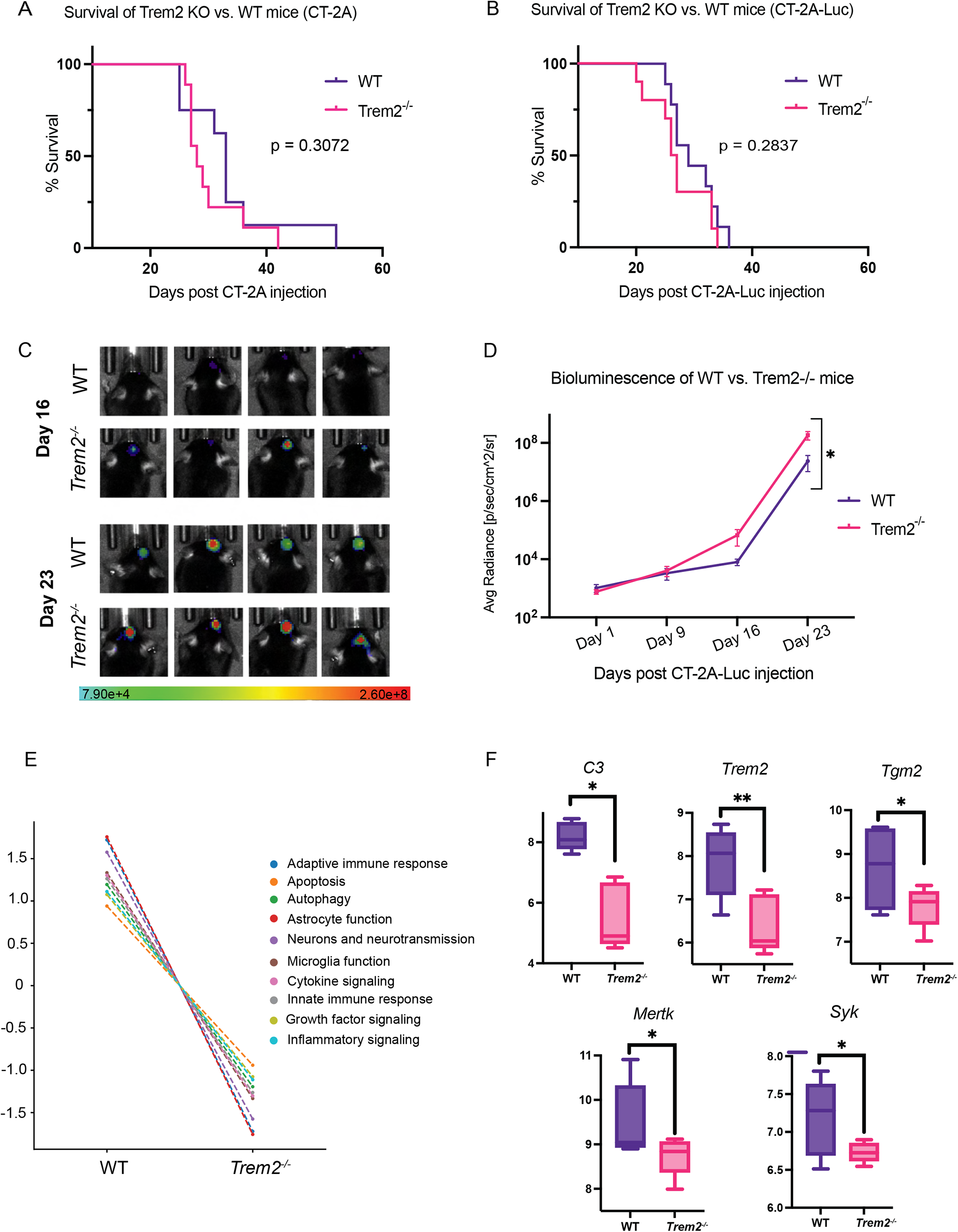
Deletion of Trem2 does not slow tumor growth in mouse models of glioma. **(A)** Kaplan-Meier curve showing survival of wildtype and *Trem2^-/-^* mice challenged with CT-2A. p=0.3072, Log-Rank (Mantel-Cox) test. **(B)** Kaplan-Meier curve of wildtype and *Trem2^-/-^* mice bearing CT-2A-luc; p = 0.2837, Log-Rank (Mantel-Cox) test. **(C)** Bioluminescence imaging of WT and *Trem2^-/-^* mice bearing CT-2A-luc. **(D)** Quantification of bioluminescence in WT and *Trem2^-/-^* mice throughout tumor progression. p=0.0135 (mixed-effects analysis). **(E)** Pathway signature scoring between WT and TREM2 KO mice. Trend plots using NanoString ncounter data were used to define pathway scores based on the expression of key genes across WT and *Trem2^-/-^* mice. Lines in the trend plot represent each pathway’s average score across WT and *Trem2^-/-^* groups. *Trem2^-/-^* mice display a drastic decrease in the pathway scores associated with adaptive immune response, apoptosis, autophagy, astrocyte function, neurons and neurotransmission, MG function, cytokine signaling, innate immune response, growth factor signaling and inflammatory signaling. **(F)** Boxplots showing the distribution of normalized log2 transformed counts from nanoString ncounter data of TREM2, C3, Tgm2, Mertk, Dock2 and Syk genes between WT and *Trem2^-/-^* mice. Median expression values of the selected genes in WT mice are higher as compared to *Trem2^-/-^* mice, and the difference is statistically significant (p-value < 0.05).

To decipher global changes that occur in the mouse brain TIME, we used the NanoString nCounter assay and measured changes in ∼700 genes involved in neuroinflammation. Brains were harvested from mice when moribund and processed to examine differential gene expression. Comparison of tumors from WT and *Trem2^-/-^* mice showed that most significantly differential genes were located in the upper left quadrant of the volcano plot implying that a majority of the genes were downregulated in the *Trem2^/-^* group (Fig. S3). To further understand the functional relevance of Trem2 in the murine TIME, we performed gene ontology analyses. Pathways that were downregulated in the *Trem2^-/-^* group compared to CT-2A tumors in the WT mice included gene families that play a role in the adaptive immune response (e.g. *Syk, Btk, Vav1*), inflammation (e.g., *Irf8, Msr1*), and MG functions (Fig. 3E, 3F and Supplementary Table S2). Of these, previous studies have linked TREM2 as anti-inflammatory modulator (Turnbull et al., 2006) and an important regulator of MG survival (Zheng et al., 2017), and it is not surprising that these functions were reduced in the TIME of *Trem2^-/-^* mice. However, several genes in the MG autophagic pathway (e.g., *Mertk, Tgm2,* and *Dock2*) were downregulated in *Trem2^-/-^* mice (Supplementary Table S2). It is noteworthy that previous studies have linked autophagic machinery to microglial phagocytosis (Berglund et al., 2020; Li et al., 2021) and TREM2 may function through these pathways to regulate glioma growth.

### TREM2^+^ MG and DC mediate phagocytosis of gliomas

To further investigate the functional role of Trem2^+^ myeloid cells in mouse gliomas, we utilized a CT-2A-ZsGreen cell line to visualize the uptake of tumor-derived fluorescent protein by myeloid cells. As ZsGreen-expressing cancer cells are phagocytosed, the fluorescence can be detected by flow cytometry, making ZsGreen a model for tracking phagocytosis and tumor antigen uptake (Bowman-Kirigin et al., 2023). We implanted CT2A-ZsGreen cells directly into the caudate nucleus of wildtype C57BL/6 mice, and 12 days after implantation, mice were sacrificed and their brains were analyzed via flow cytometry, allowing for the interrogation of Trem2^+^ZsGreen^+^ cells in MG, MAC and DCs (Fig. 4A, 4B). ZsGreen was detected in the brains of the glioma-bearing mice, specifically in myeloid subsets, confirming its efficacy as a phagocytosis surrogate. Concurrent with our human studies, we detected the presence of Trem2 on MG, MACs, and DCs (Fig. 4C). After gating on ZsGreen^+^ cells, we found that Trem2^+^ myeloid cells had higher ZsGreen expression than Trem2^-^ cells (Fig. 4D and 4E), suggesting that the phagocytic ability of Trem2^+^ cells is conserved in mice. The percent of ZsGreen^+^ cells increased for all mice in Trem2^+^ MG and DC subsets compared to Trem2^-^ MG and DCs, and the percent of ZsGreen^+^ cells was significantly higher in Trem2^+^ DC than in Trem2^-^ DC (Fig. 4F). Of note, an overwhelming majority of MACs were Trem2^+^, so a pairwise comparison between ZsGreen expression in Trem2^+^ vs. Trem2^-^ MACs was not feasible. This data indicates Trem2 promotes phagocytosis in the mouse brain TIME, and it is particularly important on DCs, suggesting a tractable potential target for enhancing phagocytosis and antigen presentation in gliomas

**Figure 4.**
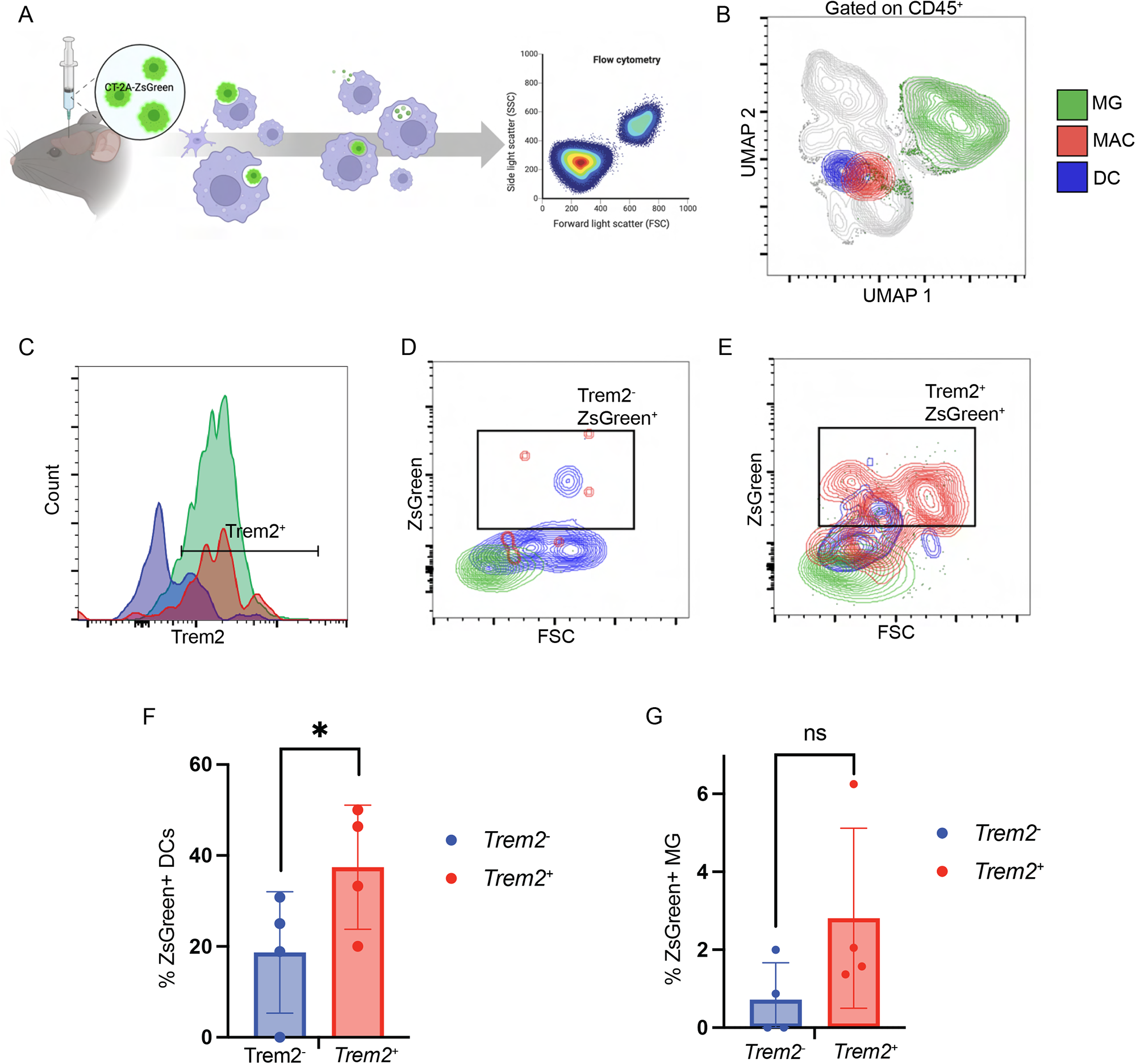
Trem2 is associated with increased ZsGreen uptake in an orthotopic murine glioma model. **(A)** Schematic outlining implantation of CT-2A-ZsGreen, phagocytosis and uptake of tumor antigen by myeloid cells, and flow cytometry analysis. Schematic illustration made in BioRender. **(B)** Uniform manifold approximation and projection (UMAP) showing MG (green), MAC (red), and DC (blue) in CT-2A-ZsGreen-bearing mouse brains. MG were defined as P2RY12^+^ CD11b^+^, MACs were defined as P2RY12^-^ CD11b^+^ F4/80^+^ MHC II^+^, and dendritic cells were defined as P2RY12^-^ CD11b^+^ CD11c^+^ MHC II^+^. **(C)** Histogram showing intensity of Trem2 expression on these distinct myeloid populations. **(D)** FSC vs. ZsGreen as gated on the Trem2^-^ population shows low proportions of Trem2^-^ZsGreen^+^ cells. **(E)** FSC vs. ZsGreen gated on Trem2^+^ populations show a large frequency of Trem2^+^ZsGreen^+^ cells. **(F)** Quantification of ZsGreen^+^ cells in Trem2^-^ vs. Trem2^+^ DCs. All mice showed higher proportions of ZsGreen in the Trem2^+^ DCs. A paired samples t test indicates that Trem2^+^/ZsGreen^+^ DCs are significantly higher in number than Trem2^-^ZsGreen^+^ DCs (p=0.0448). **(G)** Quantification of ZsGreen^+^ cells in Trem2^-^ vs. Trem2^+^ MG. A paired samples t test indicates no significant difference in Trem2^+^ZsGreen^+^ and Trem2^-^ZsGreen^+^ MG populations (p=0.2349).

Collectively, our human and mouse data position TREM2 as important regulator of phagocytosis in glioma. Emerging studies show both cell intrinsic as well immunomodulatory roles of TREM2 in cancer (Wolf et al., 2022). TREM2 is expressed in cancer cells and can function as either an oncogene or tumor suppressor (Deczkowska et al., 2020), but most studies revolve around immunosuppressive functions of TREM2 given its high expression in tumor associated MACs (Katzenelenbogen et al., 2020) (Molgora et al., 2020). As such, our studies show that blocking TREM2 in gliomas will not be feasible as an immunotherapeutic strategy as suggested in other cancers (Qiu et al., 2021). Rather, TREM2 plays a protective role like in AD, wherein loss of TREM2 alters microglial behavior including reduced phagocytosis and amyloid clearance (Keren-Shaul et al., 2017). Therefore, strategies to boost TREM2 function by stabilization and reducing its membrane shedding could be glioma specific treatment approaches that are warranted.

## Acknowledgments

This study was supported by the generous philanthropic contributions to The University of Texas (UT) MD Anderson Cancer Center (MDACC) Moon Shots Program. Part of this study was also supported by NIH grants: R21 CA222992 and R01CA225963 to K.P.B. This study was partly supported by the UT MDACC start-up research fund to L.W. We thank the UT MDACC Odyssey fellowship programs for their generous training fellowship support and the UT MDACC Divisional Research award and CPRIT pilot grant to P.G. We would like to acknowledge Lisa Norberg and Brain Tumor Center Histology Core for assistance with tissue processing, Luisa Solis Soto for assistance with IHC, Advanced Cytometry and Sorting Facility (ACSF), Small Animal Imaging Facility (SAIF) and Advanced Technology Genomics Core (ATGC) core facilities at MD Anderson for support with experiments. The ATGC facility is supported by NCI grant 520 CA016672; ACSF is supported by NCI P30 CA016672; SAIF is supported by Cancer Center Support Grant CA16672.

## Conflicts of interest

None declared.

## Data availability

Data available upon request to corresponding author.

## Author contributions

KPB, PG, and MMP conceived the study. MMP, PG, RT, SO, PC, NKM, MEM performed experiments. RT, BDV, JTH, LW, KCD assisted in data analysis. KPB and MMP wrote the manuscript with input from all authors.

---

**Supplementary Figure 1.**
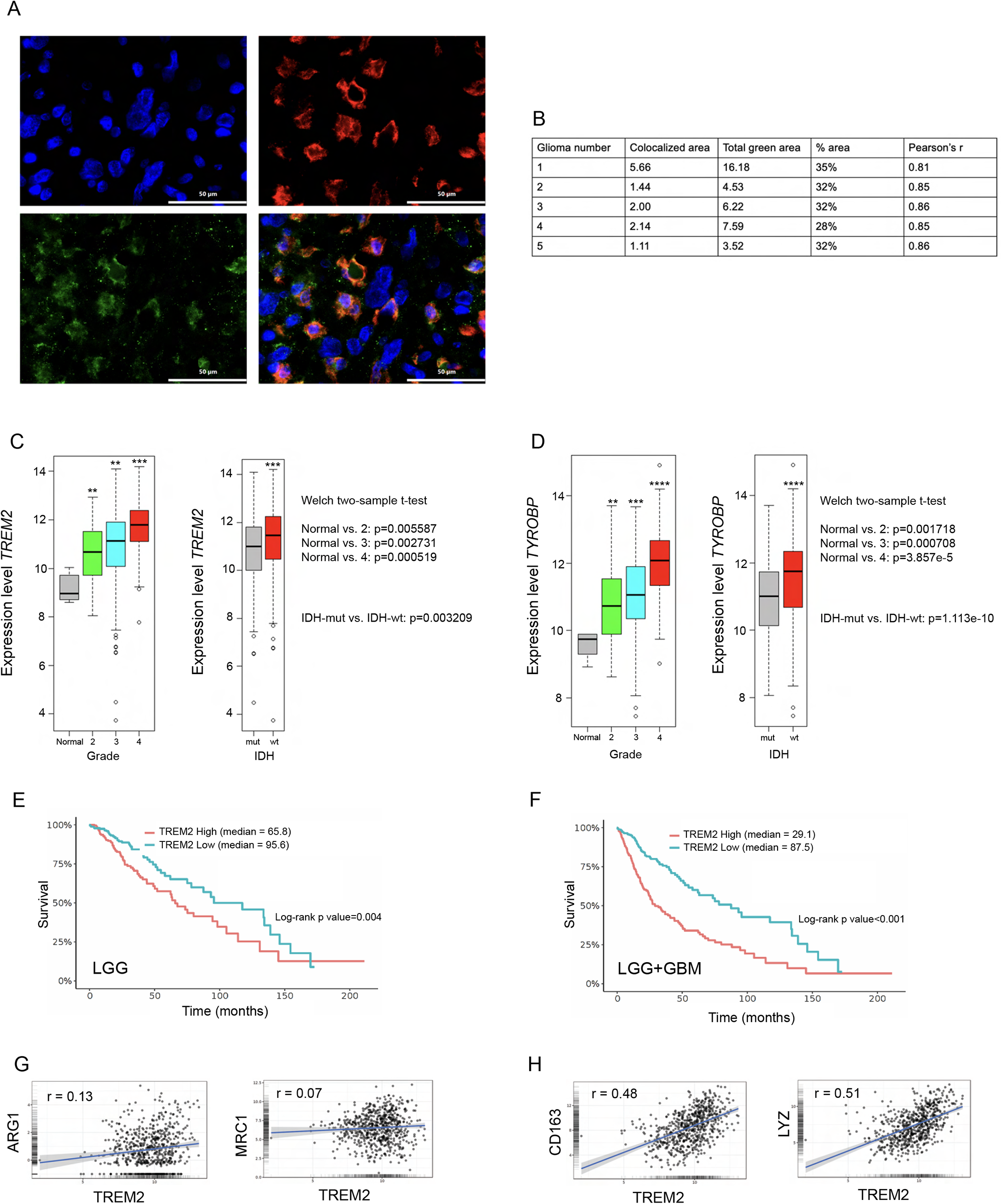
Expression of TREM2 and associated transcripts across expanded glioma subsets. **(A)** Individual and merged immunofluorescence staining images for DAPI (blue), Iba1 (red), and TREM2 (green). Scale bar: 50 µm. **(B)** Quantification of Iba1 and TREM2 colocalization in ImageJ. **(C)** Expression levels of *TREM2* from TCGA data by tumor grade, IDH classification, and subtype. **(D)** Expression levels of *TYROBP* from TCGA by tumor grade, IDH status, and subtype. **(E)** Kaplan-Meier curve for LGGs in TCGA stratified by TREM2 high or TREM2 low expression levels. Log-rank p value=0.004. **(F)** Kaplan-Meier curve for all gliomas (LGGs and GBMs pooled) in TCGA by TREM2 expression. Log-rank p value<0.00. **(G)** Linear regression plots of TREM2 vs. immunosuppression markers across all glioma types from TCGA. Pearson’s product moment correlation r = 0.13 (TREM2 vs. ARG1), p<0.001. For TREM2 vs. MRC1, r=0.07 and p=0.09. **(H)** Linear regression of TREM2 vs. phagocytic marker expression in pooled all gliomas from TCGA. Pearson’s product moment correlation for CD163 yields r=0.48 and p<0.001. For LYZ, r=0.51 and p<0.001.

**Supplementary Figure 2.**
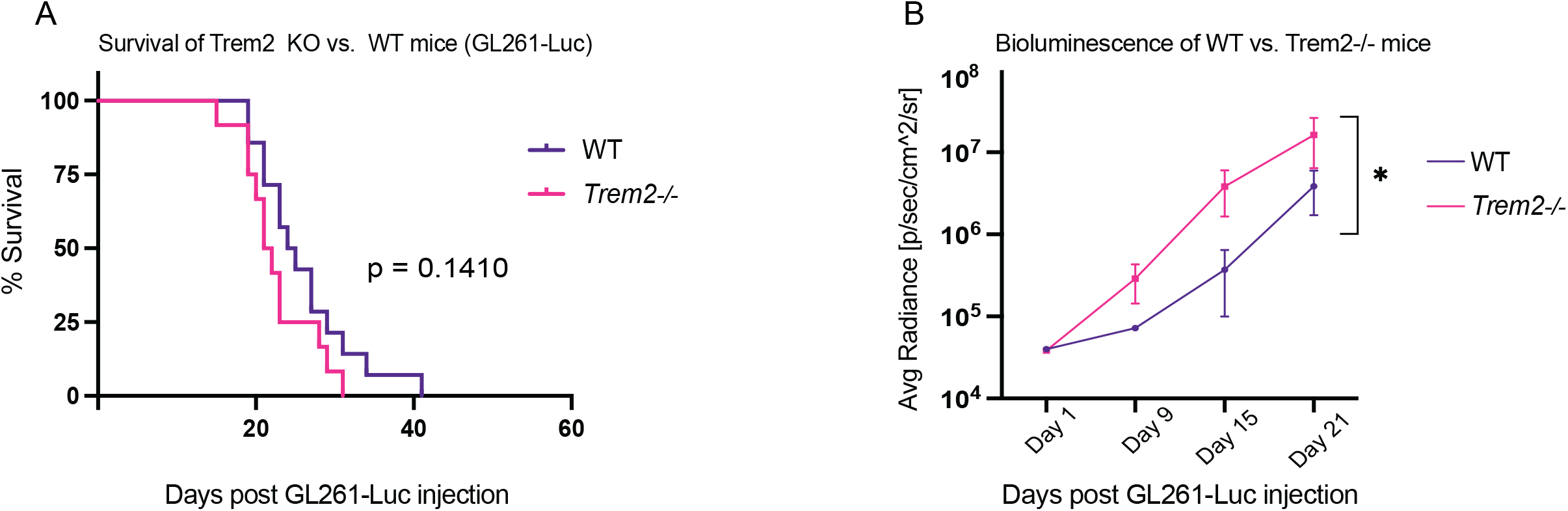
Trem2 deficiency does not improve survival in a GL261 model and is associated with increased tumor growth. **(A)** Kaplan-Meier curve showing survival differences in GL261-challenged WT and *Trem2^-/-^* mice. Log-rank p value=0.1410. **(B)** Quantification of bioluminescence of WT and *Trem2^-/-^* mice. Mixed effects analysis p=0.0158.

**Supplementary Figure 3.**
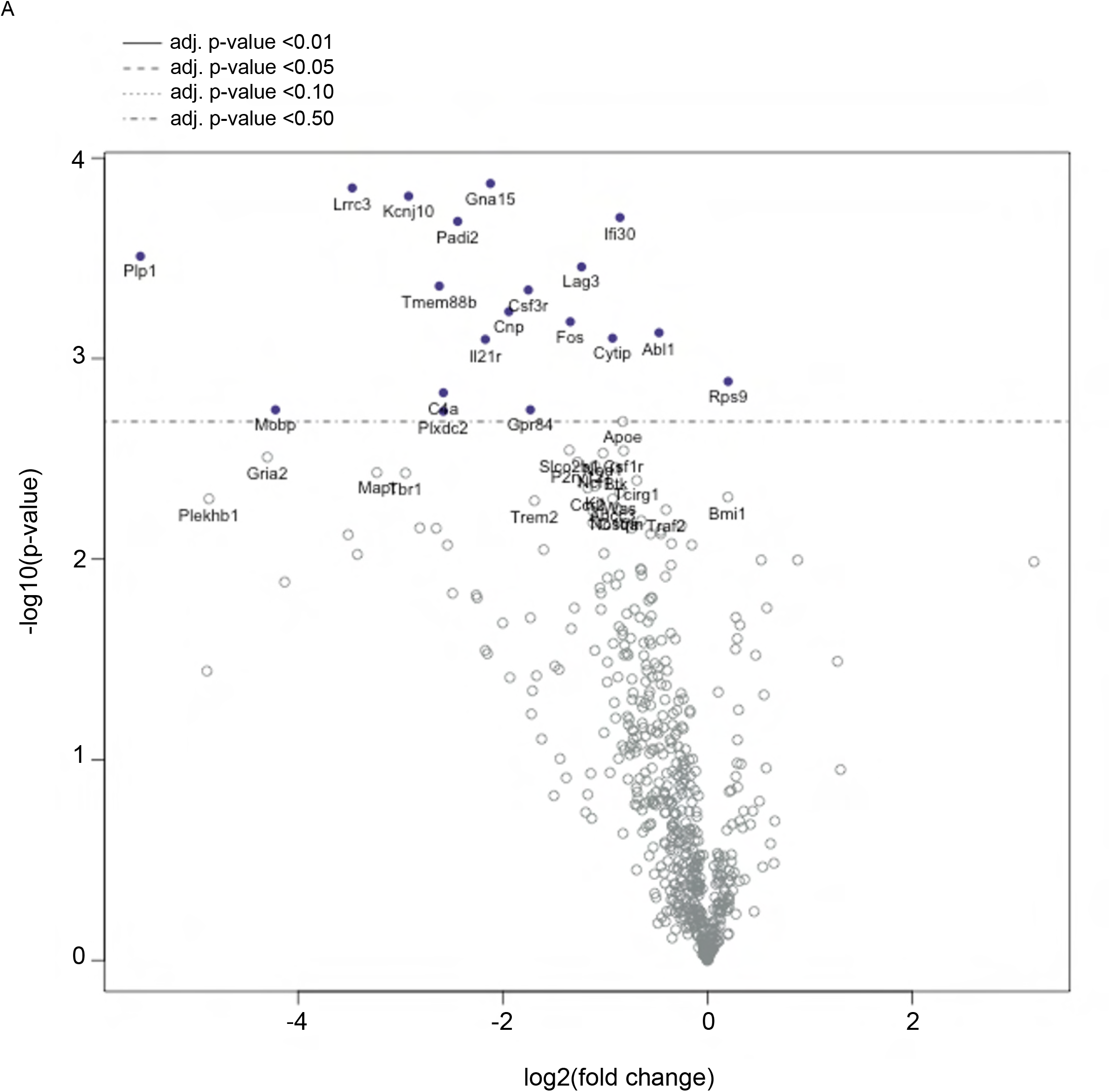
Differentially expressed genes in tumor-bearing wildtype vs. *Trem2^-/-^* mice. **(A)** Volcano plot of genes differentially expressed in NanoString analysis between WT and *Trem2^-/-^* mice. Genes left of zero (negative fold change) are downregulated in *Trem2*^-/-^ mice, and genes to the right of 0 (positive fold change) are upregulated in *Trem2^-/-^* mice compared to WT.

## Notes

### Competing Interest Statement

The authors have declared no competing interest.

## References

Barker, R.N., L.P. Erwig, K.S. Hill, A. Devine, W.P. Pearce, and A.J. Rees. 2002. Antigen presentation by macrophages is enhanced by the uptake of necrotic, but not apoptotic, cells. Clin Exp Immunol 127:220–225.

Basha, G., K. Omilusik, A. Chavez-Steenbock, A.T. Reinicke, N. Lack, K.B. Choi, and W.A. Jefferies. 2012. A CD74-dependent MHC class I endolysosomal crosspresentation pathway. Nat Immunol 13:237–245.

Berglund, R., A.O. Guerreiro-Cacais, M.Z. Adzemovic, M. Zeitelhofer, H. Lund, E. Ewing, Ruhrmann, E. Nutma, R. Parsa, M. Thessen-Hedreul, S. Amor, R.A. Harris, T. Olsson, and M. Jagodic. 2020. Microglial autophagy-associated phagocytosis is essential for recovery from neuroinflammation. Sci Immunol 5:

Bouchon, A., C. Hernandez-Munain, M. Cella, and M. Colonna. 2001. A DAP12-mediated pathway regulates expression of CC chemokine receptor 7 and maturation of human dendritic cells. J Exp Med 194:1111–1122.

Bowman, R.L., Q. Wang, A. Carro, R.G. Verhaak, and M. Squatrito. 2017. GlioVis data portal for visualization and analysis of brain tumor expression datasets. Neuro Oncol 19:139–141.

Bowman-Kirigin, J.A., R. Desai, B.T. Saunders, A.Z. Wang, M.O. Schaettler, C.J. Liu, A.J. Livingstone, D.K. Kobayashi, V. Durai, N.M. Kretzer, G.J. Zipfel, E.C. Leuthardt, J.W. Osbun, M.R. Chicoine, A.H. Kim, K.M. Murphy, T.M. Johanns, B.H. Zinselmeyer, and G.P. Dunn. 2023. The Conventional Dendritic Cell 1 Subset Primes CD8+ T Cells and Traffics Tumor Antigen to Drive Antitumor Immunity in the Brain. Cancer Immunol Res 11:20–37.

Brown, G.C., and J.J. Neher. 2014. Microglial phagocytosis of live neurons. Nat Rev Neurosci 15:209–216.

Butler, A., P. Hoffman, P. Smibert, E. Papalexi, and R. Satija. 2018. Integrating singlecell transcriptomic data across different conditions, technologies, and species. Nat Biotechnol 36:411–420.

Cancer Genome Atlas Research, N. 2008. Comprehensive genomic characterization defines human glioblastoma genes and core pathways. Nature 455:1061–1068.

Cancer Genome Atlas Research, N., D.J. Brat, R.G. Verhaak, K.D. Aldape, W.K. Yung, S.R. Salama, L.A. Cooper, E. Rheinbay, C.R. Miller, M. Vitucci, O. Morozova, A.G. Robertson, H. Noushmehr, P.W. Laird, A.D. Cherniack, R. Akbani, J.T. Huse, G. Ciriello, L.M. Poisson, J.S. Barnholtz-Sloan, M.S. Berger, C. Brennan, R.R. Colen, H. Colman, A.E. Flanders, C. Giannini, M. Grifford, A. Iavarone, R. Jain, I. Joseph, J. Kim, K. Kasaian, Mikkelsen, B.A. Murray, B.P. O’Neill, L. Pachter, D.W. Parsons, C. Sougnez, E.P. Sulman, S.R. Vandenberg, E.G. Van Meir, A. von Deimling, H. Zhang, D. Crain, K. Lau, D. Mallery, S. Morris, J. Paulauskis, R. Penny, T. Shelton, M. Sherman, P. Yena, A. Black, J. Bowen, K. Dicostanzo, J. Gastier-Foster, K.M. Leraas, T.M. Lichtenberg, C.R. Pierson, N.C. Ramirez, C. Taylor, S. Weaver, L. Wise, E. Zmuda, T. Davidsen, J.A. Demchok, G. Eley, M.L. Ferguson, C.M. Hutter, K.R. Mills Shaw, B.A. Ozenberger, M. Sheth, H.J. Sofia, R. Tarnuzzer, Z. Wang, L. Yang, J.C. Zenklusen, B. Ayala, J. Baboud, S. Chudamani, M.A. Jensen, J. Liu, T. Pihl, R. Raman, Y. Wan, Y. Wu, A. Ally, J.T. Auman, M. Balasundaram, S. Balu, S.B. Baylin, R. Beroukhim, M.S. Bootwalla, R. Bowlby, C.A. Bristow, D. Brooks, Y. Butterfield, R. Carlsen, S. Carter, L. Chin, A. Chu, E. Chuah, K. Cibulskis, A. Clarke, S.G. Coetzee, N. Dhalla, T. Fennell, S. Fisher, S. Gabriel, G. Getz, R. Gibbs, R. Guin, A. Hadjipanayis, D.N. Hayes, T. Hinoue, K. Hoadley, R.A. Holt, A.P. Hoyle, S.R. Jefferys, S. Jones, C.D. Jones, R. Kucherlapati, P.H. Lai, E. Lander, S. Lee, L. Lichtenstein, Y. Ma, D.T. Maglinte, H.S. Mahadeshwar, M.A. Marra, M. Mayo, S. Meng, M.L. Meyerson, P.A. Mieczkowski, R.A. Moore, L.E. Mose, A.J. Mungall, A. Pantazi, M. Parfenov, P.J. Park, J.S. Parker, C.M. Perou, A. Protopopov, X. Ren, J. Roach, T.S. Sabedot, J. Schein, S.E. Schumacher, J.G. Seidman, S. Seth, H. Shen, J.V. Simons, P. Sipahimalani, M.G. Soloway, X. Song, H. Sun, B. Tabak, A. Tam, D. Tan, J. Tang, N. Thiessen, T. Triche, Jr., D.J. Van Den Berg, U. Veluvolu, S. Waring, D.J. Weisenberger, M.D. Wilkerson, T. Wong, J. Wu, L. Xi, A.W. Xu, L. Yang, T.I. Zack, J. Zhang, B.A. Aksoy, H. Arachchi, C. Benz, B. Bernard, D. Carlin, J. Cho, D. DiCara, S. Frazer, G.N. Fuller, J. Gao, N. Gehlenborg, D. Haussler, D.I. Heiman, L. Iype, A. Jacobsen, Z. Ju, S. Katzman, H. Kim, T. Knijnenburg, R.B. Kreisberg, M.S. Lawrence, W. Lee, K. Leinonen, P. Lin, S. Ling, W. Liu, Y. Liu, Y. Liu, Y. Lu, G. Mills, S. Ng, M.S. Noble, E. Paull, A. Rao, S. Reynolds, G. Saksena, Z. Sanborn, C. Sander, N. Schultz, Y. Senbabaoglu, R. Shen, I. Shmulevich, R. Sinha, J. Stuart, S.O. Sumer, Y. Sun, N. Tasman, B.S. Taylor, D. Voet, N. Weinhold, J.N. Weinstein, D. Yang, K. Yoshihara, S. Zheng, W. Zhang, L. Zou, T. Abel, S. Sadeghi, M.L. Cohen, J. Eschbacher, E.M. Hattab, A. Raghunathan, M.J. Schniederjan, D. Aziz, G. Barnett, W. Barrett, D.D. Bigner, L. Boice, C. Brewer, C. Calatozzolo, B. Campos, C.G. Carlotti, Jr., T.A. Chan, L. Cuppini, E. Curley, S. Cuzzubbo, K. Devine, F. DiMeco, R. Duell, J.B. Elder, A. Fehrenbach, G. Finocchiaro, W. Friedman, J. Fulop, J. Gardner, B. Hermes, C. Herold-Mende, C. Jungk, A. Kendler, N.L. Lehman, E. Lipp, O. Liu, R. Mandt, M. McGraw, R. McLendon, C. McPherson, L. Neder, P. Nguyen, A. Noss, R. Nunziata, Q.T. Ostrom, C. Palmer, A. Perin, B. Pollo, A. Potapov, O. Potapova, W.K. Rathmell, D. Rotin, L. Scarpace, C. Schilero, K. Senecal, K. Shimmel, V. Shurkhay, S. Sifri, R. Singh, A.E. Sloan, K. Smolenski, S.M. Staugaitis, R. Steele, L. Thorne, D.P. Tirapelli, A. Unterberg, M. Vallurupalli, Y. Wang, R. Warnick, F. Williams, Y. Wolinsky, S. Bell, M. Rosenberg, C. Stewart, F. Huang, J.L. Grimsby, A.J. Radenbaugh, and J. Zhang. 2015. Comprehensive, Integrative Genomic Analysis of Diffuse Lower-Grade Gliomas. N Engl J Med 372:2481–2498.

Cassetta, L., and J.W. Pollard. 2018. Targeting macrophages: therapeutic approaches in cancer. Nat Rev Drug Discov 17:887–904.

Ceccarelli, M., F.P. Barthel, T.M. Malta, T.S. Sabedot, S.R. Salama, B.A. Murray, O. Morozova, Y. Newton, A. Radenbaugh, S.M. Pagnotta, S. Anjum, J. Wang, G. Manyam, P. Zoppoli, S. Ling, A.A. Rao, M. Grifford, A.D. Cherniack, H. Zhang, L. Poisson, C.G. Carlotti, Jr., D.P. Tirapelli, A. Rao, T. Mikkelsen, C.C. Lau, W.K. Yung, R. Rabadan, J. Huse, D.J. Brat, N.L. Lehman, J.S. Barnholtz-Sloan, S. Zheng, K. Hess, G. Rao, M. Meyerson, R. Beroukhim, L. Cooper, R. Akbani, M. Wrensch, D. Haussler, K.D. Aldape, P.W. Laird, D.H. Gutmann, T.R. Network, H. Noushmehr, A. Iavarone, and R.G. Verhaak. 2016. Molecular Profiling Reveals Biologically Discrete Subsets and Pathways of Progression in Diffuse Glioma. Cell 164:550–563.

Chen, Z., X. Feng, C.J. Herting, V.A. Garcia, K. Nie, W.W. Pong, R. Rasmussen, B. Dwivedi, S. Seby, S.A. Wolf, D.H. Gutmann, and D. Hambardzumyan. 2017. Cellular and Molecular Identity of Tumor-Associated Macrophages in Glioblastoma. Cancer Res 77:2266–2278.

Colton, C., and D.M. Wilcock. 2010. Assessing activation states in microglia. CNS Neurol Disord Drug Targets 9:174–191.

Consortium, G. 2018. Glioma through the looking GLASS: molecular evolution of diffuse gliomas and the Glioma Longitudinal Analysis Consortium. Neuro Oncol 20:873884.

Deczkowska, A., A. Weiner, and I. Amit. 2020. The Physiology, Pathology, and Potential Therapeutic Applications of the TREM2 Signaling Pathway. Cell 181:1207–1217.

Friebel, E., K. Kapolou, S. Unger, N.G. Nunez, S. Utz, E.J. Rushing, L. Regli, M. Weller, M. Greter, S. Tugues, M.C. Neidert, and B. Becher. 2020. Single-Cell Mapping of Human Brain Cancer Reveals Tumor-Specific Instruction of Tissue-Invading Leukocytes. Cell 181:1626–1642 e1620.

Gabrusiewicz, K., B. Rodriguez, J. Wei, Y. Hashimoto, L.M. Healy, S.N. Maiti, G. Thomas, S. Zhou, Q. Wang, A. Elakkad, B.D. Liebelt, N.K. Yaghi, R. Ezhilarasan, N. Huang, J.S. Weinberg, S.S. Prabhu, G. Rao, R. Sawaya, L.A. Langford, J.M. Bruner, G.N. Fuller, A. Bar-Or, W. Li, R.R. Colen, M.A. Curran, K.P. Bhat, J.P. Antel, L.J. Cooper, E.P. Sulman, and A.B. Heimberger. 2016. Glioblastoma-infiltrated innate immune cells resemble M0 macrophage phenotype. JCI Insight 1:

Guerreiro, R., A. Wojtas, J. Bras, M. Carrasquillo, E. Rogaeva, E. Majounie, C. Cruchaga, C. Sassi, J.S. Kauwe, S. Younkin, L. Hazrati, J. Collinge, J. Pocock, T. Lashley, J. Williams, J.C. Lambert, P. Amouyel, A. Goate, R. Rademakers, K. Morgan, J. Powell, P. St George-Hyslop, A. Singleton, J. Hardy, and G. Alzheimer Genetic Analysis. 2013. TREM2 variants in Alzheimer’s disease. N Engl J Med 368:117127.

Hegi, M.E., A.C. Diserens, T. Gorlia, M.F. Hamou, N. de Tribolet, M. Weller, J.M. Kros, J.A. Hainfellner, W. Mason, L. Mariani, J.E. Bromberg, P. Hau, R.O. Mirimanoff, J.G. Cairncross, R.C. Janzer, and R. Stupp. 2005. MGMT gene silencing and benefit from temozolomide in glioblastoma. N Engl J Med 352:997–1003.

Hutter, G., J. Theruvath, C.M. Graef, M. Zhang, M.K. Schoen, E.M. Manz, M.L. Bennett, A. Olson, T.D. Azad, R. Sinha, C. Chan, S. Assad Kahn, S. Gholamin, C. Wilson, G. Grant, J. He, I.L. Weissman, S.S. Mitra, and S.H. Cheshier. 2019. Microglia are effector cells of CD47-SIRPalpha antiphagocytic axis disruption against glioblastoma. Proc Natl Acad Sci U S A 116:997–1006.

Jonsson, T., H. Stefansson, S. Steinberg, I. Jonsdottir, P.V. Jonsson, J. Snaedal, S. Bjornsson, J. Huttenlocher, A.I. Levey, J.J. Lah, D. Rujescu, H. Hampel, I. Giegling, O.A. Andreassen, K. Engedal, I. Ulstein, S. Djurovic, C. Ibrahim-Verbaas, A. Hofman, M.A. Ikram, C.M. van Duijn, U. Thorsteinsdottir, A. Kong, and K. Stefansson. 2013. Variant of TREM2 associated with the risk of Alzheimer’s disease. N Engl J Med 368:107–116.

Katzenelenbogen, Y., F. Sheban, A. Yalin, I. Yofe, D. Svetlichnyy, D.A. Jaitin, C. Bornstein, A. Moshe, H. Keren-Shaul, M. Cohen, S.Y. Wang, B. Li, E. David, T.M. Salame, A. Weiner, and I. Amit. 2020. Coupled scRNA-Seq and Intracellular Protein Activity Reveal an Immunosuppressive Role of TREM2 in Cancer. Cell 182:872–885 e819.

Keren-Shaul, H., A. Spinrad, A. Weiner, O. Matcovitch-Natan, R. Dvir-Szternfeld, T.K. Ulland, E. David, K. Baruch, D. Lara-Astaiso, B. Toth, S. Itzkovitz, M. Colonna, M. Schwartz, and I. Amit. 2017. A Unique Microglia Type Associated with Restricting Development of Alzheimer’s Disease. Cell 169:1276–1290 e1217.

Khantakova, D., S. Brioschi, and M. Molgora. 2022. Exploring the Impact of TREM2 in Tumor-Associated Macrophages. Vaccines (Basel) 10:

Kim, W.J., Y.S. Dho, C.Y. Ock, J.W. Kim, S.H. Choi, S.T. Lee, I.H. Kim, T.M. Kim, and C.K. Park. 2019. Clinical observation of lymphopenia in patients with newly diagnosed glioblastoma. J Neurooncol 143:321–328.

Kleinberger, G., M. Brendel, E. Mracsko, B. Wefers, L. Groeneweg, X. Xiang, C. Focke, M. Deussing, M. Suarez-Calvet, F. Mazaheri, S. Parhizkar, N. Pettkus, W. Wurst, R. Feederle, P. Bartenstein, T. Mueggler, T. Arzberger, I. Knuesel, A. Rominger, and C. Haass. 2017. The FTD-like syndrome causing TREM2 T66M mutation impairs microglia function, brain perfusion, and glucose metabolism. EMBO J 36:1837–1853.

Klemm, F., R.R. Maas, R.L. Bowman, M. Kornete, K. Soukup, S. Nassiri, J.P. Brouland, C.A. Iacobuzio-Donahue, C. Brennan, V. Tabar, P.H. Gutin, R.T. Daniel, M.E. Hegi, and J.A. Joyce. 2020. Interrogation of the Microenvironmental Landscape in Brain Tumors Reveals Disease-Specific Alterations of Immune Cells. Cell 181:1643–1660 e1617.

Krasemann, S., C. Madore, R. Cialic, C. Baufeld, N. Calcagno, R. El Fatimy, L. Beckers, E. O’Loughlin, Y. Xu, Z. Fanek, D.J. Greco, S.T. Smith, G. Tweet, Z. Humulock, T. Zrzavy, P. Conde-Sanroman, M. Gacias, Z. Weng, H. Chen, E. Tjon, F. Mazaheri, K. Hartmann, A. Madi, J.D. Ulrich, M. Glatzel, A. Worthmann, J. Heeren, B. Budnik, C. Lemere, T. Ikezu, F.L. Heppner, V. Litvak, D.M. Holtzman, H. Lassmann, H.L. Weiner, J. Ochando, C. Haass, and O. Butovsky. 2017. The TREM2-APOE Pathway Drives the Transcriptional Phenotype of Dysfunctional Microglia in Neurodegenerative Diseases. Immunity 47:566–581 e569.

Lal, S., M. Lacroix, P. Tofilon, G.N. Fuller, R. Sawaya, and F.F. Lang. 2000. An implantable guide-screw system for brain tumor studies in small animals. J Neurosurg 92:326–333.

Larkin, C.J., V.A. Arrieta, H. Najem, G. Li, P. Zhang, J. Miska, P. Chen, C.D. James, A.M. Sonabend, and A.B. Heimberger. 2022. Myeloid Cell Classification and Therapeutic Opportunities Within the Glioblastoma Tumor Microenvironment in the Single Cell-Omics Era. Front Immunol 13:907605.

Li, G., P. Sherchan, Z. Tang, and J. Tang. 2021. Autophagy & Phagocytosis in Neurological Disorders and their Possible Cross-talk. Curr Neuropharmacol 19:1912–1924.

Molgora, M., E. Esaulova, W. Vermi, J. Hou, Y. Chen, J. Luo, S. Brioschi, M. Bugatti, A.S. Omodei, B. Ricci, C. Fronick, S.K. Panda, Y. Takeuchi, M.M. Gubin, R. Faccio, M. Cella, S. Gilfillan, E.R. Unanue, M.N. Artyomov, R.D. Schreiber, and M. Colonna. 2020. TREM2 Modulation Remodels the Tumor Myeloid Landscape Enhancing Anti-PD-1 Immunotherapy. Cell 182:886–900 e817.

Morantz, R.A., G.W. Wood, M. Foster, M. Clark, and K. Gollahon. 1979. Macrophages in experimental and human brain tumors. Part 1: Studies of the macrophage content of experimental rat brain tumors of varying immunogenicity. J Neurosurg 50:298304.

N’Diaye, E.N., C.S. Branda, S.S. Branda, L. Nevarez, M. Colonna, C. Lowell, J.A. Hamerman, and W.E. Seaman. 2009. TREM-2 (triggering receptor expressed on myeloid cells 2) is a phagocytic receptor for bacteria. J Cell Biol 184:215–223.

Nimmerjahn, A., F. Kirchhoff, and F. Helmchen. 2005. Resting microglial cells are highly dynamic surveillants of brain parenchyma in vivo. Science 308:1314–1318.

Peng, Q., S. Malhotra, J.A. Torchia, W.G. Kerr, K.M. Coggeshall, and M.B. Humphrey. 2010. TREM2- and DAP12-dependent activation of PI3K requires DAP10 and is inhibited by SHIP1. Sci Signal 3:ra38.

Podetz-Pedersen, K.M., V. Vezys, N.V. Somia, S.J. Russell, and R.S. McIvor. 2014. Cellular immune response against firefly luciferase after sleeping beauty-mediated gene transfer in vivo. Hum Gene Ther 25:955–965.

Pyonteck, S.M., L. Akkari, A.J. Schuhmacher, R.L. Bowman, L. Sevenich, D.F. Quail, O.C. Olson, M.L. Quick, J.T. Huse, V. Teijeiro, M. Setty, C.S. Leslie, Y. Oei, A. Pedraza, J. Zhang, C.W. Brennan, J.C. Sutton, E.C. Holland, D. Daniel, and J.A. Joyce. 2013. CSF-1R inhibition alters macrophage polarization and blocks glioma progression. Nat Med 19:1264–1272.

Qiu, H., Z. Shao, X. Wen, J. Jiang, Q. Ma, Y. Wang, L. Huang, X. Ding, and L. Zhang. 2021. TREM2: Keeping Pace With Immune Checkpoint Inhibitors in Cancer Immunotherapy. Front Immunol 12:716710.

Quail, D.F., and J.A. Joyce. 2017. The Microenvironmental Landscape of Brain Tumors. Cancer Cell 31:326–341.

Reardon, D.A., A.A. Brandes, A. Omuro, P. Mulholland, M. Lim, A. Wick, J. Baehring, M.S. Ahluwalia, P. Roth, O. Bahr, S. Phuphanich, J.M. Sepulveda, P. De Souza, S. Sahebjam, M. Carleton, K. Tatsuoka, C. Taitt, R. Zwirtes, J. Sampson, and M. Weller. 2020. Effect of Nivolumab vs Bevacizumab in Patients With Recurrent Glioblastoma: The CheckMate 143 Phase 3 Randomized Clinical Trial. JAMA Oncol 6:1003–1010.

Sade-Feldman, M., K. Yizhak, S.L. Bjorgaard, J.P. Ray, C.G. de Boer, R.W. Jenkins, D.J. Lieb, J.H. Chen, D.T. Frederick, M. Barzily-Rokni, S.S. Freeman, A. Reuben, P.J. Hoover, A.C. Villani, E. Ivanova, A. Portell, P.H. Lizotte, A.R. Aref, J.P. Eliane, M.R. Hammond, H. Vitzthum, S.M. Blackmon, B. Li, V. Gopalakrishnan, S.M. Reddy, Z.A. Cooper, C.P. Paweletz, D.A. Barbie, A. Stemmer-Rachamimov, K.T. Flaherty, J.A. Wargo, G.M. Boland, R.J. Sullivan, G. Getz, and N. Hacohen. 2018. Defining T Cell States Associated with Response to Checkpoint Immunotherapy in Melanoma. Cell 175:998–1013 e1020.

Sanchez, V.E., J.P. Lynes, S. Walbridge, X. Wang, N.A. Edwards, A.K. Nwankwo, H.P. Sur, G.A. Dominah, A. Obungu, N. Adamstein, P.K. Dagur, D. Maric, J. Munasinghe, J.D. Heiss, and E.K. Nduom. 2020. GL261 luciferase-expressing cells elicit an anti-tumor immune response: an evaluation of murine glioma models. Sci Rep 10:11003.

Sasaki, Y., K. Ohsawa, H. Kanazawa, S. Kohsaka, and Y. Imai. 2001. Iba1 is an actincross-linking protein in macrophages/microglia. Biochem Biophys Res Commun 286:292–297.

Schetters, S.T.T., D. Gomez-Nicola, J.J. Garcia-Vallejo, and Y. Van Kooyk. 2017. Neuroinflammation: Microglia and T Cells Get Ready to Tango. Front Immunol 8:1905.

Schulz, D., Y. Severin, V.R.T. Zanotelli, and B. Bodenmiller. 2019. In-Depth Characterization of Monocyte-Derived Macrophages using a Mass CytometryBased Phagocytosis Assay. Sci Rep 9:1925.

Stupp, R., W.P. Mason, M.J. van den Bent, M. Weller, B. Fisher, M.J. Taphoorn, K. Belanger, A.A. Brandes, C. Marosi, U. Bogdahn, J. Curschmann, R.C. Janzer, S.K. Ludwin, T. Gorlia, A. Allgeier, D. Lacombe, J.G. Cairncross, E. Eisenhauer, R.O. Mirimanoff, R. European Organisation for, T. Treatment of Cancer Brain, G. Radiotherapy, and G. National Cancer Institute of Canada Clinical Trials. 2005. Radiotherapy plus concomitant and adjuvant temozolomide for glioblastoma. N Engl J Med 352:987–996.

Takahashi, K., C.D. Rochford, and H. Neumann. 2005. Clearance of apoptotic neurons without inflammation by microglial triggering receptor expressed on myeloid cells2. J Exp Med 201:647–657.

Timperi, E., P. Gueguen, M. Molgora, I. Magagna, Y. Kieffer, S. Lopez-Lastra, P. Sirven, L.G. Baudrin, S. Baulande, A. Nicolas, G. Champenois, D. Meseure, A. VincentSalomon, A. Tardivon, E. Laas, V. Soumelis, M. Colonna, F. Mechta-Grigoriou, S. Amigorena, and E. Romano. 2022. Lipid-Associated Macrophages Are Induced by Cancer-Associated Fibroblasts and Mediate Immune Suppression in Breast Cancer. Cancer Res 82:3291–3306.

Turnbull, I.R., S. Gilfillan, M. Cella, T. Aoshi, M. Miller, L. Piccio, M. Hernandez, and M. Colonna. 2006. Cutting edge: TREM-2 attenuates macrophage activation. J Immunol 177:3520–3524.

Ulland, T.K., W.M. Song, S.C. Huang, J.D. Ulrich, A. Sergushichev, W.L. Beatty, A.A. Loboda, Y. Zhou, N.J. Cairns, A. Kambal, E. Loginicheva, S. Gilfillan, M. Cella, H.W. Virgin, E.R. Unanue, Y. Wang, M.N. Artyomov, D.M. Holtzman, and M. Colonna. 2017. TREM2 Maintains Microglial Metabolic Fitness in Alzheimer’s Disease. Cell 170:649–663 e613.

Ulrich, J.D., M.B. Finn, Y. Wang, A. Shen, T.E. Mahan, H. Jiang, F.R. Stewart, L. Piccio, M. Colonna, and D.M. Holtzman. 2014. Altered microglial response to Abeta plaques in APPPS1-21 mice heterozygous for TREM2. Mol Neurodegener 9:20.

Venezie, R.D., A.D. Toews, and P. Morell. 1995. Macrophage recruitment in different models of nerve injury: lysozyme as a marker for active phagocytosis. J Neurosci Res 40:99–107.

Wang, Y., M. Cella, K. Mallinson, J.D. Ulrich, K.L. Young, M.L. Robinette, S. Gilfillan, G.M. Krishnan, S. Sudhakar, B.H. Zinselmeyer, D.M. Holtzman, J.R. Cirrito, and M. Colonna. 2015. TREM2 lipid sensing sustains the microglial response in an Alzheimer’s disease model. Cell 160:1061–1071.

Wang, Y., T.K. Ulland, J.D. Ulrich, W. Song, J.A. Tzaferis, J.T. Hole, P. Yuan, T.E. Mahan, Y. Shi, S. Gilfillan, M. Cella, J. Grutzendler, R.B. DeMattos, J.R. Cirrito, D.M. Holtzman, and M. Colonna. 2016. TREM2-mediated early microglial response limits diffusion and toxicity of amyloid plaques. J Exp Med 213:667–675.

Wei, J., P. Chen, P. Gupta, M. Ott, D. Zamler, C. Kassab, K.P. Bhat, M.A. Curran, J.F. de Groot, and A.B. Heimberger. 2020. Immune biology of glioma-associated macrophages and microglia: functional and therapeutic implications. Neuro Oncol 22:180–194.

Wolf, E.M., B. Fingleton, and A.H. Hasty. 2022. The therapeutic potential of TREM2 in cancer. Front Oncol 12:984193.

Yeh, F.L., Y. Wang, I. Tom, L.C. Gonzalez, and M. Sheng. 2016. TREM2 Binds to Apolipoproteins, Including APOE and CLU/APOJ, and Thereby Facilitates Uptake of Amyloid-Beta by Microglia. Neuron 91:328–340.

Yu, M., Y. Chang, Y. Zhai, B. Pang, P. Wang, G. Li, T. Jiang, and F. Zeng. 2022. TREM2 is associated with tumor immunity and implies poor prognosis in glioma. Front Immunol 13:1089266.

Zheng, H., L. Jia, C.C. Liu, Z. Rong, L. Zhong, L. Yang, X.F. Chen, J.D. Fryer, X. Wang, Y.W. Zhang, H. Xu, and G. Bu. 2017. TREM2 Promotes Microglial Survival by Activating Wnt/beta-Catenin Pathway. J Neurosci 37:1772–1784.

